# Stochastic Gene Expression in Proliferating Cells: Differing Noise Intensity in Single-Cell and Population Perspectives

**DOI:** 10.1101/2024.06.28.601263

**Authors:** Zhanhao Zhang, Iryna Zabaikina, César Nieto, Zahra Vahdat, Pavol Bokes, Abhyudai Singh

## Abstract

Random fluctuations (noise) in gene expression can be studied from two complementary perspectives: following expression in a single cell over time or comparing expression between cells in a proliferating population at a given time. Here, we systematically investigated scenarios where both perspectives lead to different levels of noise in a given gene product. We first consider a stable protein, whose concentration is diluted by cellular growth, and the protein inhibits growth at high concentrations, establishing a positive feedback loop. For a stochastic model with molecular bursting of gene products, we analytically predict and contrast the steady-state distributions of protein concentration in both frameworks. Although positive feedback amplifies the noise in expression, this amplification is much higher in the population framework compared to following a single cell over time. We also study other processes that lead to different noise levels even in the absence of such dilution-based feedback. When considering randomness in the partitioning of molecules between daughters during mitosis, we find that in the single-cell perspective, the noise in protein concentration is independent of noise in the cell cycle duration. In contrast, partitioning noise is amplified in the population perspective by increasing randomness in cell-cycle time. Overall, our results show that the commonly used single-cell framework that does not account for proliferating cells can, in some cases, underestimate the noise in gene product levels. These results have important implications for studying the inter-cellular variation of different stress-related expression programs across cell types that are known to inhibit cellular growth.

## Introduction

The intracellular level of gene products is the result of complex interconnected biochemical processes that are intrinsically stochastic and often operate with low-copy number components. This stochasticity is manifested as intercellular variation in gene expression levels within an isogenic cell population despite controlling for factors, such as the extracellular environment and cell-cycle effects [1–4]. Random fluctuations (noise) in gene expression levels fundamentally impact all aspects of cell physiology and the fidelity of cellular information processing. Not surprisingly, depending on the gene function and context, expression noise is subject to evolutionary pressures [5–10] and actively regulated through diverse mechanisms. For example, the promoter architecture/genomic environment [11–13], the kinetics of different gene expression steps [14–16], the inclusion of feedback/feedforward loops [17–22], and the ubiquitous binding of proteins to *decoy sites* [23–25] have been shown to both attenuate or amplify noise levels.

Over the last few decades, single-cell studies have interestingly revealed the beneficial roles of noise in gene product levels. These include, but are not limited to, driving genetically-identical cells to different fates [26–33] and facilitating population adaptation to environmental fluctuations [34–38]. The latter scenario is exemplified by rare populations of clonal cells that survive lethal stresses, as seen in antibiotic treatment of bacteria [39–44], or cancer cells undergoing chemotherapy [45–49]. The non-genetic basis of heterogeneous single-cell responses to stress is a topic of current research, and several publications have implicated pre-existing drug-tolerant expression states arising as result of noise in gene regulatory networks [50, 51].

Random fluctuations in the level of a given protein can be studied from two perspectives: single cell and population [52–54]. The commonly used single-cell perspective approach captures the stochastic dynamics of protein level in a single cell over time (Figure 1A), and here the effects of cell growth and division are either ignored or implicitly captured (for example, through the continuous dilution of concentration). In the population perspective, one explicitly considers an exponentially expanding cell population, and gene product variability is quantified across all cells at a given time point. A fundamental question of interest is *when do these complementary perspectives predict different degrees of stochastic variation in gene expression?*

**Figure 1:**
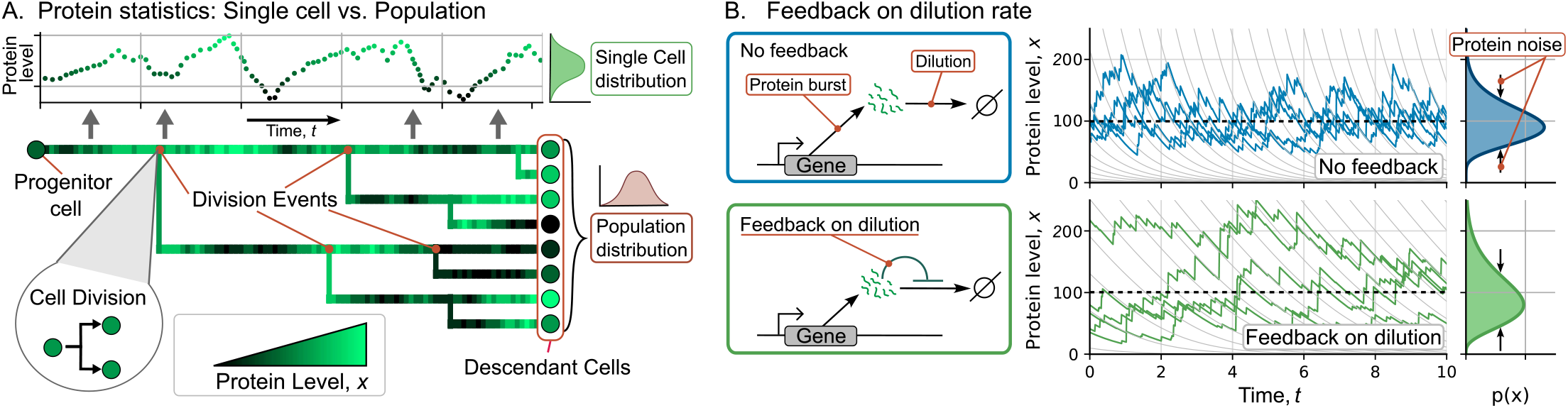
Single-cell and population perspectives for investigating stochastic gene expression with dilution-based feedback regulation. (**A**) *(top:)* For the single-cell perspective, concentration of a given protein is tracked along a single lineage. *(bottom:)* From a population point of view, the protein concentration distribution is obtained across all descendants of the colony. Different shades of green represent the protein level for each cell. (**B**) *(left:)* Schematic of the gene expression model with random bursts of protein synthesis, and concentration dilution in between burst events. In the model without regulation (blue), the dilution rate is constant. In the model with feedback on dilution (green), the dilution rate decreases as the protein concentration increases, according to (3). *(right:)* Sample trajectories and corresponding protein concentration distributions for both models. The gray lines in the background show protein dilution trajectories; horizontal dashed lines represent the mean concentration in both models. Parameter values used for these trajectories are *β* = 10, *k* = 1*/*100, *λ* = 10 (no feedback), *λ* = 4.76 (with feedback), *γ* = 1 in arbitrary units. The time axis is normalized with respect to the dilution rate *γ*.

Our analysis identifies two scenarios where single-cell and population perspectives yield different extents of fluctuations in the concentration of a given protein of interest. *The first scenario arises when the intracellular concentration directly or indirectly affects cellular growth*, and hence determines the cell’s proliferation capacity. We specifically focus on the case where high protein concentration inhibits cellular growth. This drives the concentration even higher because of reduced dilution. This effect implements a positive feedback loop [55–58]. This expression-growth coupling can be seen in many cases of protein-induced stress response [59], cell resource saturation [60, 61], and is a feature of many stress-tolerant expression programs. For example, high expression of specific proteins comes at the cost of inhibiting cellular growth in the absence of stress, but improves cell survival in the presence of stress [62–65].

From a mathematical perspective, we propose approaches based on the solution of the associated differential Chapman-Kolmogorov equation (dCKE), and the population balance equation (PBE), to derive protein concentration distributions in the single-cell and population perspectives, respectively. The simplicity of our modeling frameworks allows exact derivations of the corresponding probability density function (pdf), which are then compared and contrasted between the two perspectives with increasing feedback strength.

*The second scenario corresponds to randomness in the partitioning of protein molecules between two daughters during mitosis and cytokinesis* [66]. In this case, concentration fluctuations in the single-cell perspective are modeled using the formalism of Stochastic Hybrid Systems (SHS) resulting in an exact analytical formula for the concentration noise level, as quantified by the steady-state squared coefficient of variation of protein concentration. The corresponding statistics from the population perspective are obtained via agent-based models that track expression levels within each cell of a proliferating colony. Interestingly, our results show that coupling the partitioning process with the inherent randomness of cell-cycle times enhances expression variability across the population as compared to the single-cell perspective. We begin by formally introducing the two different perspectives for studying stochastic expression, and how stochasticity is modeled based on random bursts of gene activity.

### Single cell and population to quantify the statistics of gene expression

We study stochastic variations in the concentration of a specific protein within proliferating cells using two complementary perspectives. These perspectives are graphically illustrated in Figure 1A, where the expansion of a cell colony is represented as a lineage tree: the root of the tree represents the progenitor cell, each branching point is a cell division event, and the horizontal distance between consecutive branching points represents the cell-cycle duration. The color intensity represents the protein level at a given time: the lighter the green, the higher the protein level. During cell division, a mother cell splits into two identical daughters, each inheriting half of the mother’s volume and protein amount. Thus, the protein concentration in the newborn daughters is assumed to be equal to the mother’s concentration just before division. This assumption of perfect partitioning will be relaxed later in the manuscript. In the single-cell approach, only one of the two daughter cells is tracked after division and protein statistics are determined based on a single lineage path over time. In contrast, in the population approach, both daughter cells are tracked, and statistics are estimated on all descendant cells at a fixed point in time.

To analytically derive and contrast the concentration distribution in both perspectives, we take advantage of a simple one-dimensional model of gene expression that has previously been introduced [67] and validated with single-cell data [68, 69]. This model consists of protein synthesis that occurs in short periods of intense gene activity, often referred to in the literature as *bursting*. Diverse mechanisms that span all stages of gene expression (promoter activation / transcription / translation) have been attributed to bursting [70–77], and each stochastically-occurring burst event increases protein concentration by a random amount. Considering a long-lived protein (i.e., half-life much longer than the cell-doubling time), its concentration is continuously diluted along the cell cycle due to exponential growth in cell size.

### Effects of growth-mediated feedbacks on protein concentration fluctuations

In this section, we provide descriptions of the model coupling bursting expression events with dilution-based positive feedback in both single-cell and population perspectives. These models are analyzed to derive *exact* steady-state distributions for protein concentration that are compared and contrasted between the two perspectives with and without feedback regulation.

#### Protein distribution for single-cell perspective

In the single-cell perspective, the protein concentration *x* (*t*) at time *t* within an individual cell evolves stochastically according to the following rules. Burst events occur according to a Poisson process with rate or burst frequency *λ*. During each *burst, x* increases instantly with a burst size *b* drawn from an exponential distribution with mean *β*. These increments in concentration are conveniently represented by the reset:

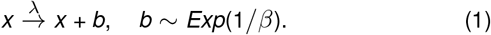

It is important to point out that both the burst frequency and size in this concentration model are invariant with respect to the cell size. This implicitly assumes appropriate scaling of expression rates (in terms of the number of molecules synthesised per unit time) with cell size [78–86].

Feedback in dilution is modeled phenomenologically by considering the cellular growth rate

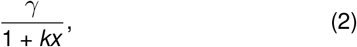

to be a decreasing function of concentration *x* (Figure 1B, lower panels). This results in the following dilution dynamics in between burst events

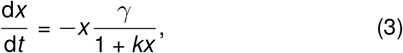

where *k* ≥ 0 can be interpreted as the *feedback strength* and *γ* > 0 is the *maximum dilution rate*. In summary, intracellular fluctuations in protein concentration are captured by the piecewise deterministic Markov process (PDMP) *x* (*t*) defined by (1)-(3). For the reader’s convenience, we provide a compilation of model parameters and symbols used to quantify concentration statistics in Table 1.

**Table 1:**
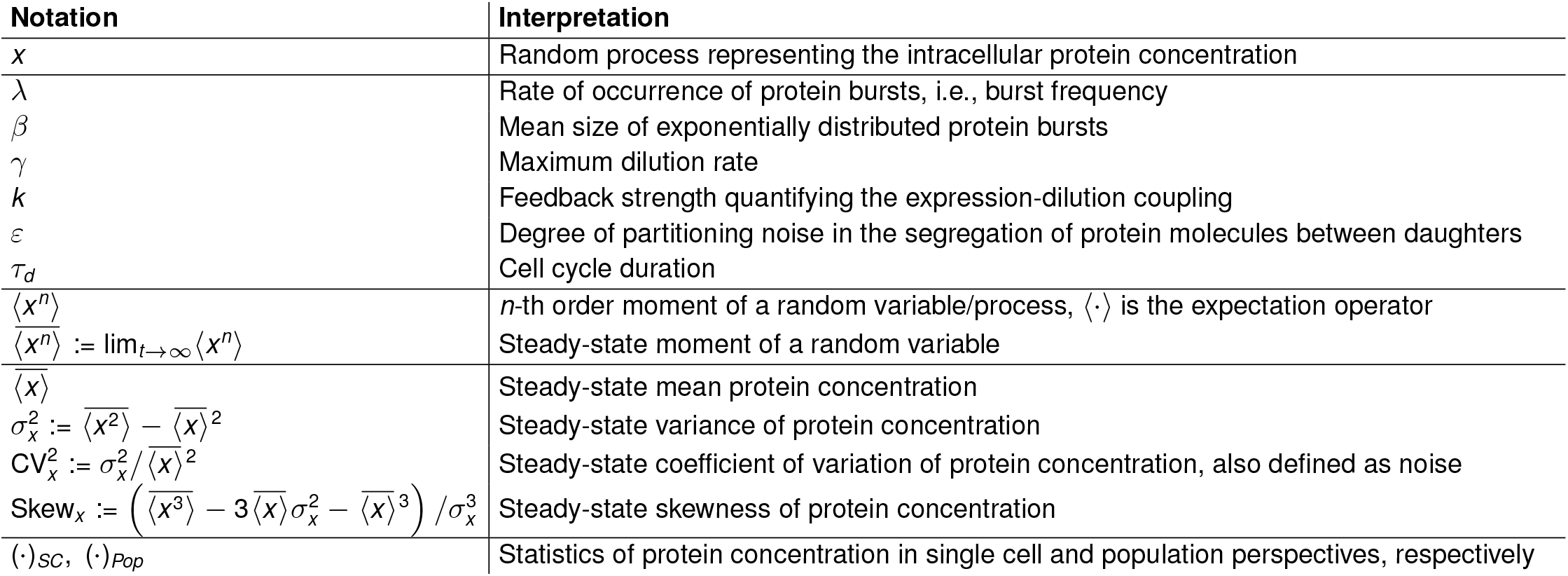
Model parameters and variables studied in the text.

Before analyzing the feedback model, we briefly review the special case of no feedback (*k* = 0) that corresponds to constant cellular growth and dilution rate *γ* (Figure 1B, upper panels). Prior analysis of this unregulated gene expression model predicted that the steady-state protein distribution *p*(*x*) followed a gamma distribution with parameters *λ/γ* and *β* (shape and scale, respectively) [67], consistent with single-cell variations in the expression of specific proteins in *Escherichia coli* and *Saccsharomyces cerevisiae* [69]. The steady-state protein distribution can be analyzed through its statistical properties (defined in Table 1): the mean 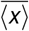, the squared coefficient of variation 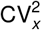 (quantifying the noise in protein level), and the skewness (measure of the distribution asymmetry). For this unregulated case (*k* = 0), we obtain

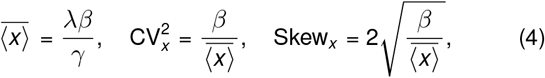

and it is interesting to note the ratio Skew_*x*_ */*CV_*x*_ = 2.

Now, considering the model with feedback (*k >* 0), the time evolution of the probability density function (pdf) *p*(*x, t*)_*SC*_ is given by the differential Chapman-Kolmogorov equation (dCKe) [67, 87, 88]. Here and throughout the article the subscript *SC* is used to denote distributions and statistics in the single-cell perspective. The stationary protein distribution *p*(*x*)_*SC*_ defined as *p*(*x*)_*SC*_ := lim_*t→∞*_ *p*_*SC*_ (*x, t*) follows

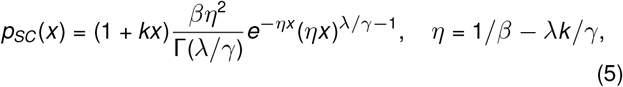

where 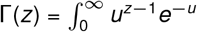 is the gamma function (see Appendix S1 for details on dCKe and its analytical solution). We observe from (5) that *p*_*SC*_ (*x*) exists only for the set of parameters *λ, β* and *k* that satisfy *η >* 0. An interpretation of this condition is that the average production flow *λβ* must be less than the maximum dilution flow *γ/k* in (3). Otherwise, the protein dilution is not fast enough to compensate for the protein production rate and the mean level unboundedly increases over time; thus, the stationary distribution does not exist. Figure 2A shows the space of parameters where the distribution (5) exists.

**Figure 2:**
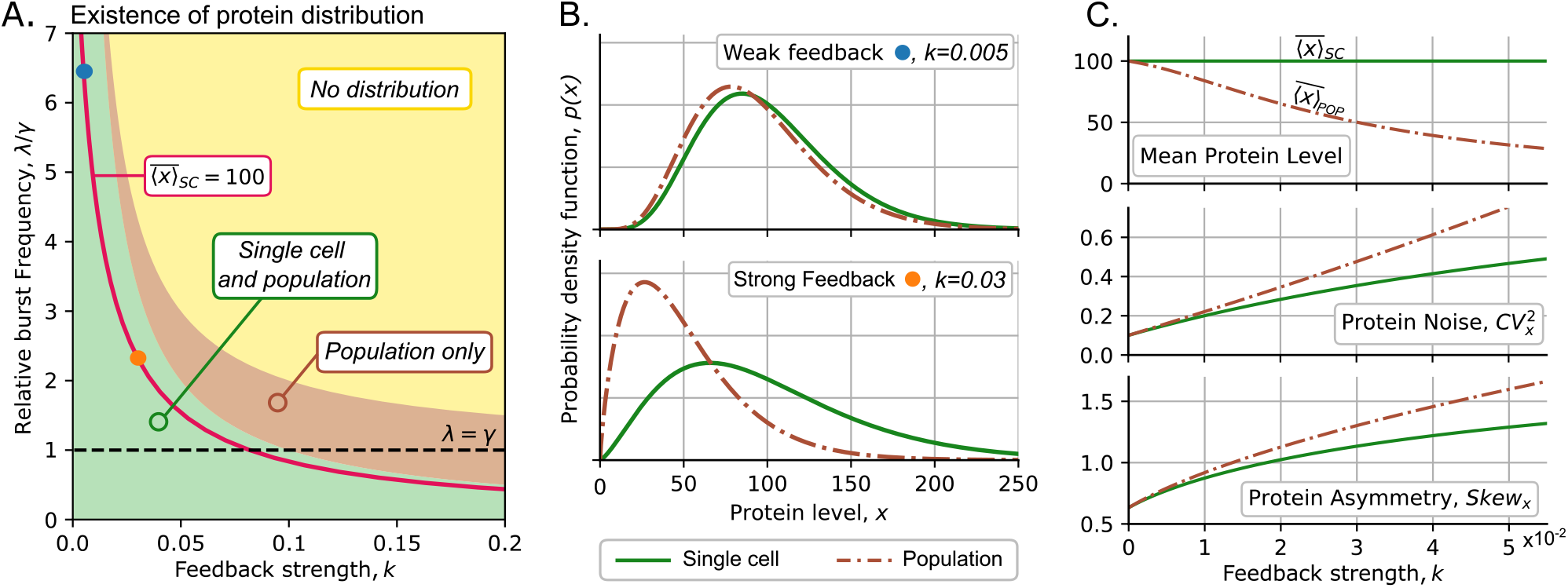
Comparison of protein distribution in single-cell and population perspectives as the feedback intensifies. (**A**) The region of existence of the steady-state protein distribution in terms of the feedback strength *k* and relative burst frequency *λ/γ. Green region*: stationary distribution *p*(*x*) exists in both single cell and population perspectives. *Brown region*: Only the distribution in the population perspective exists. *Yellow region*: Distribution does not exist in any of the frameworks. *Bold red line*: a set of values (*k, λ/γ*), resulting in fixed 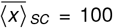 as per (7). (**B**) Comparison of protein distribution in single-cell (solid green line) and population perspectives (brown dashed) as feedback increases: *(top:)* weak feedback, *(bottom:)* strong feedback. (**C**) *(from top to bottom:)* Mean protein level, protein noise, and distribution asymmetry in the single cell and population frameworks; *λ* is chosen so that 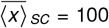 as *k* increases following the bold red line in panel (**A**). For all plots, we set *β* = 10, *γ* = 1.

Using (5) we obtain the following statistical properties of the protein level in terms of the feedback strength *k*

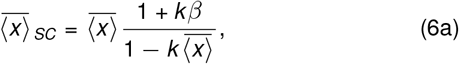

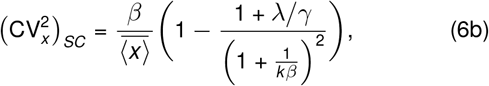

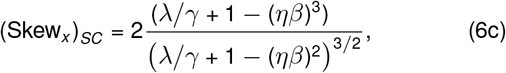

where 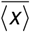 is the mean concentration of the unregulated process as per (4). To have a fair comparison of the effect of increasing *k*, we hold the mean protein level 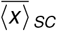 fixed for different values of *k*. To achieve this, we use (6a) to express the burst frequency *λ* as a function of *k* :

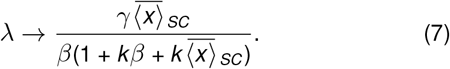

As a graphic example, the way *λ* must change as we increase *k* to maintain 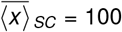 is shown in Figure 2A (red line). Figure 2C shows that after fixing 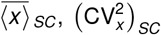, and (Skew_*x*_)_*SC*_ are increasing functions of *k* (green lines). Thus, as feedback becomes stronger, the protein distribution exhibits greater noise and becomes more right-skewed. In particular, in the limit of weak feedback strength (*k* ≪1), the distribution statistics are approximated as:

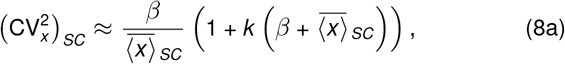

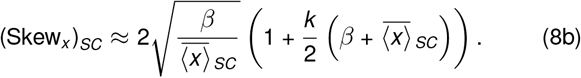

Notice that without regulation (*k* = 0), the protein distribution in a single cell becomes identical to the unregulated one with statistics given by (4). The presence of weak regulation scales these unregulated statistics (4) by an increasing function of *k*. Finally, recall that the ratio ((Skew_*x*_)_*SC*_ */* (CV_*x*_)_*SC*_) was equal to 2 for unregulated gene expression (*k* = 0), but decreases below 2 with increasing positive feedback strength *k*.

#### Protein distribution for population perspective

To extend the single-cell framework to a population one, it is required to describe the dynamics of cell proliferation. We assume that cell division events are modeled by a non-homogeneous Poisson process with rate *γ/*(1 + *kx*) [89]. Then a cell with the protein level *x* at time *t* has a probability [*γ/*(1 + *kx*)]*dt* to divide during the next infinitesimal time interval (*t, t* + *dt*). It also follows that cells with low protein concentrations proliferate faster than those with high protein levels. Note that in the limit of the unregulated protein (*k* = 0), division events occur according to the standard Poisson process with rate *γ*, and the cell cycle is exponentially distributed with mean 1*/γ*.

In the population framework, in addition to the protein, we also quantify the time evolution of the cell population size. We introduce the population density function *h*(*x, t*), which describes the population as the number of cells with a given concentration *x* at time *t*. Then *h*(*x, t*) is obtained by solving the population balance equation (PBE) [54, 90]. The PBE is similar to the dCKE, but after division, the process follows the dynamics of both daughter cells. In steady-state conditions, only the population size grows, whereas the proportion of cells with a given protein level remains steady. Thus, *h*(*x, t*) can be decomposed as

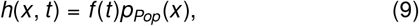

where *f* (*t*) is an exponential function (explained in Appendix S2) associated with the population size growth and *p*_*Pop*_(*x*) is the stationary probability density function of the protein concentration *x*; the subscript *Pop* represents quantities determined in the population perspective. In Appendix S2, we also show that *p*_*Pop*_(*x*) has the closed expression:

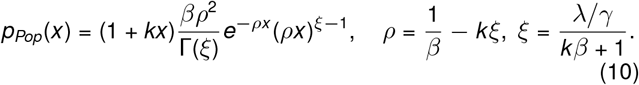

Similarly to the single-cell approach, the protein distribution at population perspective exists if *ρ >* 0. Figure 2A shows that the population distribution exists whenever the single-cell distribution does. Moreover, *p*_*Pop*_(*x*) always exists when *λ < γ*, that is, when the burst frequency is below the maximum dilution rate. In general, the population distribution exists for values of *λ* that satisfy (*λ*−*γ*)*β < γ/k*. This existence condition is less strict than the one for a single-cell model: a population distribution may exist even if the single-cell distribution does not (Figure 2A).

A comparison of the protein distributions for both frameworks is shown in Figure 2B. If the feedback is weak (upper panel), then the difference between the distributions is insignificant; this is consistent with the fact that both distributions are identical to the gamma distribution in the unregulated case (*k* = 0). As the feedback becomes stronger (lower panel), the differences between protein distributions become larger: population distribution shifts to lower concentration and becomes more light-tailed, indicating a larger portion of cells with low protein concentration.

The statistics of the protein level in the population perspective are obtained as:

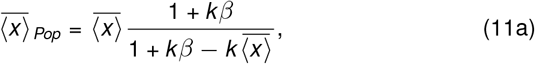

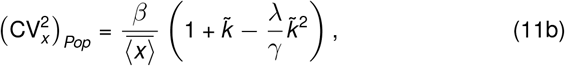

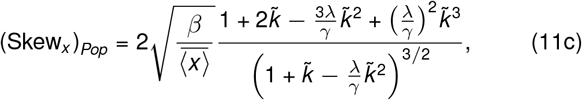

where 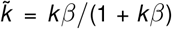 is an auxiliary constant, and these are compared to their counterparts in the single-cell perspectives in Figure 2C. To obtain the limit of weak feedback strength (*k*≪ 1), we take the approach used in (8) and fix 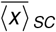. The statistics for the protein from population perspective (11) can be expressed in terms of their single-cell counterparts and the parameter *k* :

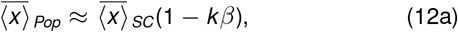

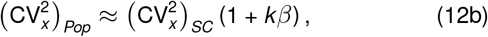

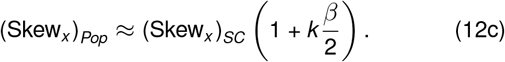

We see that from the population perspective, noise and skewness (given by (12b) and (12c), respectively) increase at least linearly faster than the moments for a single cell. This is because as the feedback intensifies, the population includes more fast-proliferating cells with low protein concentration, and the mean protein level 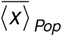 decreases to zero, as shown in Figure 2C (upper panel). This causes higher noise levels for stronger feedback (Figure 2C, middle panel). Finally, as *k* increases, the population distribution becomes more right-skewed (Figure 2C, lower panel) as a result of population composition shifting to proliferative cells with lower concentrations.

We proceed with limit of strong feedback, as we keep 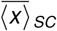 fixed according to (7). We observe that the statistics in each perspective show different properties. Noise and skewness in the single-cell framework exhibit the following limits:

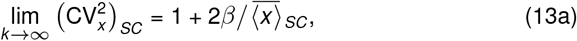

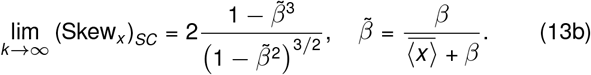

In contrast, these statistics are unbounded in the population perspective, i.e.,

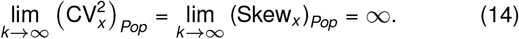

In a single cell, the existence conditions provide an exact counterbalancing relationship between the feedback strength and the intrinsic noise (stochastic production); thus, the noise is finite. In the population perspective, the existence condition is relaxed, so the feedback does not fully counterbalance the intrinsic noise.

As a final remark, we also performed an analysis of how the protein statistics change when we increase the feedback strength *k* and keep parameters *λ, β*, and *γ* fixed. Under these conditions we obtained similar results: noise level and skewness in single cell perspective are always lower than in the population one; further details and comparison to unregulated case are provided in Appendix S3.

Now, having explored the impact of growth-mediated feedback on protein statistics, we study how additional noise generated during the partitioning of protein molecules between daughter cells impacts protein stochasticity in single-cell and population perspectives.

### Effects of molecule partitioning on protein concentration fluctuations

During cell division or mitosis, a parent cell segregates its contents, including chromosomes, organelles, and gene products, between daughters. In the absence of any active regulation of segregation, each molecule has a random chance of being inherited by each descendant cell. This randomness in partitioning constitutes an additional noise source that drives intercellular heterogeneity in protein levels [66, 91–95]. We investigated whether this source of noise makes different predictions on expression variability in single-cell and population perspectives. To isolate the effect of partitioning noise, we first explore an unregulated model in which gene expression evolves deterministically with no intrinsic noise. Next, we relax this assumption by incorporating stochastic bursting events.

#### Coupling deterministic expression with partitioning errors

As a starting point, we ignore noise in gene expression and the protein concentration *x* evolves deterministically as per the first-order differential equation:

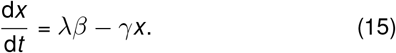

The term *λβ* (the product of the burst frequency and the average burst size) represents the net average protein synthesis rate and *γ* is the constant dilution rate ignoring feedback regulation.

In the previous section, we assume that the cell cycle duration, the time between consecutive divisions, is exponentially distributed. Relaxing this assumption, we now consider the cell-cycle time to be an independent and identically distributed random variable *τ*_*d*_ that can follow any arbitrary positively-valued continuous distribution (Figure 3A). We set the mean duration of the cell cycle as ⟨*τ*_*d*_⟩ = ln 2*/γ* and quantify the noise in *τ*_*d*_ through its squared coefficient of variation, 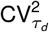.

**Figure 3:**
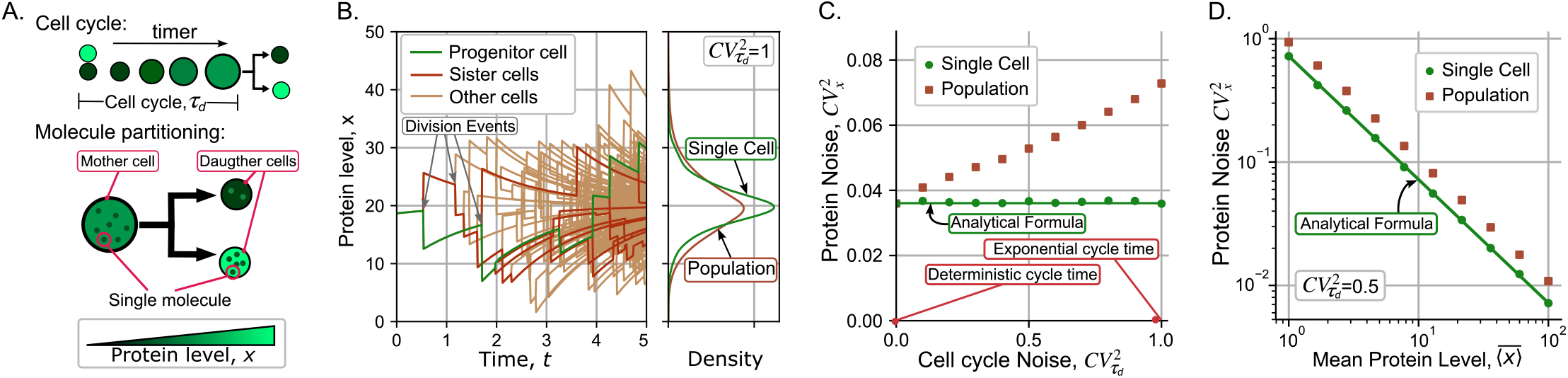
Noisy cell-cycle durations amplify protein noise differences between single-cell and population perspectives. (**A**) *(top:)* The cell-cycle duration *τ*_*d*_ is a random variable following an arbitrary distribution. Within the cell cycle, the protein concentration evolves deterministically as per the ordinary differential equation (15). *(bottom:)* During mitosis, protein molecules are randomly segregated among daughters resulting in differences in the inherited concentration. Different shades of green represent different levels of protein concentration. (**B**) *(left:)* Trajectories of protein concentration in an expanding cell colony, where jumps represent randomness in protein partitioning among daughters during cell division. The green line: a single-cell trajectory is generated by randomly choosing one of the two daughter cells (red lines) after each division event. The brown lines represent other descendant cells. The cell-cycle times are assumed to be exponentially distributed in this simulation. *(right:)* The steady-state probability density functions of the protein concentration in single-cell and population perspectives. (**C**) Effect of noise in the cell cycle time as quantified by its squared coefficient of variation 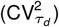 on the noise in the protein concentration 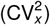. The solid line is the analytically predicted noise in the single-cell perspective as given by (18), and the dots represent noise levels computed from simulations. Mean concentrations in both models are identical 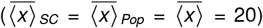. (**D**) A logarithmic scale representation of the steady-state protein noise level as a function of the mean protein level, highlighting variability differences between single-cell and population perspectives. Parameters used for the plot are 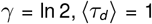, *ε* = 1, *kβ* = 20*γ*. Population statistics were calculated using cells of 2000 colonies and 5000 individuals were simulated for the single cell perspective. Statistics were calculated after 6 generations.

Having defined the timing of cell-division events, we next describe how we model the random partitioning during the process. First, consider a mother cell with concentration *x* just before division. A division event results in two daughters with concentrations *x*^+^ and 2*x*−*x*^+^ respectively. Here *x*^+^ is a random variable that is appropriately bounded 0 *< x*^+^ *<* 2*x* to ensure non-negative concentrations, and has the following mean and variance (conditioned on *x*)

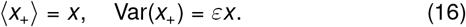

Note that the mean protein concentration in both newborn daughters is the same as the mother cell. The constant *ε* quantifies the extent of randomness in the partitioning process and depends on multiple of factors, such as the cell size at mitosis, the specifics of molecular segregation (for example, molecules segregating as monomers or dimers), errors in cell size partitioning, etc. Note that the scenario of perfect partitioning (*x*_+_ = *x* with probability one) is recovered for *ε* = 0. We refer the reader to Appendix S4 for further details on this approach and how *x*^+^ is randomly generated.

#### Partitioning noise for single-cell perspective

In the single-cell perspective, *x* (*t*) is a PDMP with deterministic dynamics (15) and resets

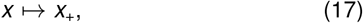

that occur during the division events (Figure 3B). If the protein level after the division *x*^+^ follows the statistics (16), it is possible to obtain exact analytical formulas for the steady-state mean and noise levels of *x* (*t*) (see Appendix S5 for details). More specifically, our results show that for any arbitrarily distributed cell-cycle time *τ*_*d*_, the protein statistics are

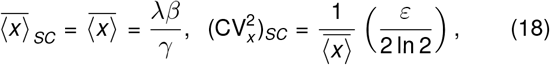

and remarkably, they are invariant of the statistical properties of the cell-cycle time. Thus, making the cell-cycle times more random, this is, increasing 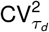 for fixed mean ⟨*τ*_*d*_ ⟩, will not have any impact on the protein noise level (Figure 3C). Notice the inverse scaling of noise 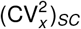 with the mean protein level in (18) that is a direct consequence of the variance of *x*_+_ being proportional to the concentration in (16). This is also seen in the unregulated bursting model (4) leading to indistinguishability of noise mechanisms from such scaling relationships [66].

#### Partitioning noise for population perspective

Having analytically solved the statistics of concentration fluctuations in the single-cell perspective, we turn our attention to quantifying protein variability in an expanding cell colony. In Appendix S6 we show that a fixed cell-cycle duration (this is *τ*_*d*_ = ⟨ *τ*_*d*_⟩ with probability one) yields the same concentration noise in both perspectives.

For general randomly distributed cell-cycle duration, we resort to simulation of agent-based models as done in previous works [96, 97]. The basis of the algorithm, with more details in Appendix S7 and published in our repository [98], consists of considering each cell as an agent with particular properties, such as protein level *x* and an internal timer that regulates its division timing. Each division event leads to two newborn cells with concentration partitioning as described above. During the cell cycle, protein concentrations evolve by (15) considering deterministic expression, and the time to the next division is drawn independently according to a prescribed arbitrary statistical distribution.

Sample realizations of protein concentrations from this agent-based framework are illustrated in Figure 3B. Both perspectives yield the same mean concentration while the protein level noise follows:

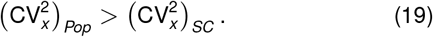

This means that protein has a higher noise in protein concentration from the population perspective. A quantification of this difference is presented in Figure 3C, where both noise levels start out equal when 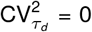. In contrast to the single-cell perspective where 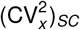 is invariant of 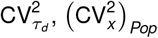 increases monotonically with increasing randomness in cell-cycle duration. For exponentially-distributed cell-cycle times 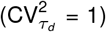, the approximation used by several models [99], and as assumed in the PBE model of the previous section, 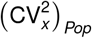 is approximately twice of 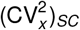 (Figure 3C). The inverse scaling of noise with mean as seen in (18) is also preserved in the population perspective, albeit with a higher proportionality constant resulting in a shifted line in the logarithmic scale (Figure 3D) (similar to that seen in (12b) for the case of dilution-based feedback).

Finally, the qualitative differences seen in Figure 3C are recapitulated in more realistic agent-based models that explicitly take into account cell size dynamics (see Figure S3 in Appendix S8), and here cell size homeostasis mechanisms drive mother-daughter and daughter-daughter cell-cycle correlations.

#### Coupling stochastic expression with partitioning errors

To complete the approach, we now consider stochastic gene expression as captured by protein synthesis occurring in random bursts. From a single-cell perspective, the protein noise is elegantly derived as (see details in Appendix S5)

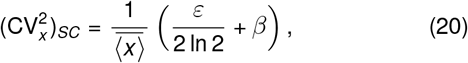

and is the sum of two noise contributions as given by equations (4) (contribution form intrinsic noise) and (18) (contribution form partitioning noise). To obtain the corresponding statistics in the population perspective we modify the earlier described agent-based model to consider intracellular concentrations evolving via stochastic bursts and continuous exponential decay with a constant dilution rate *γ* in between bursts. Figure 4 shows simulation trajectories corrupting to two different scenarios:

**Figure 4:**
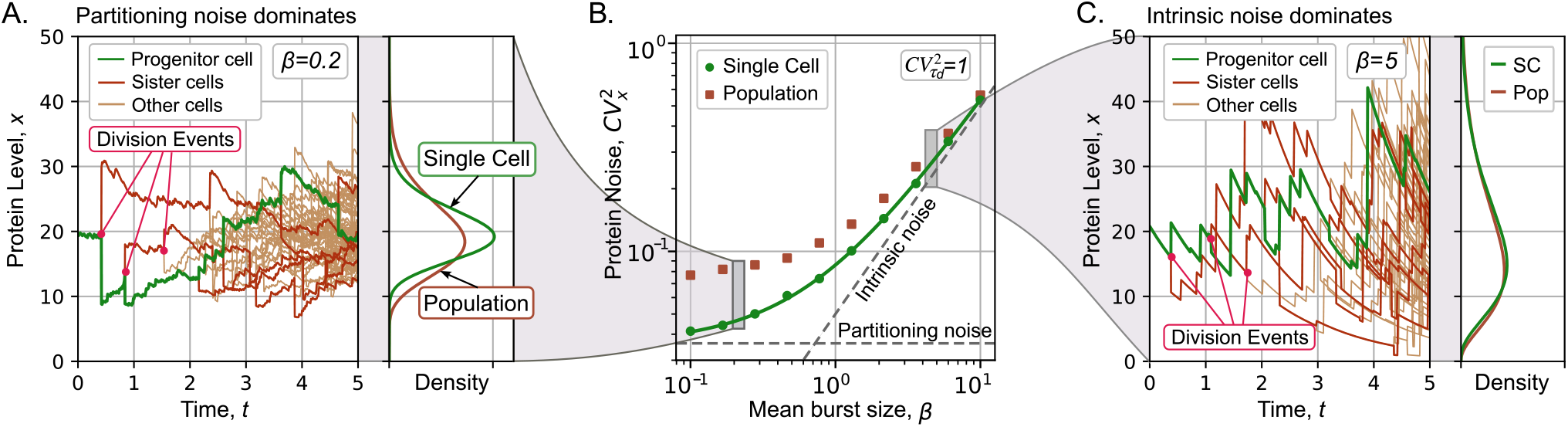
Increasing randomness in molecular segregation between daughters enhances protein noise differences between single-cell and population perspectives. (**A**) Sample trajectories of protein concentration in an expanding cell colony when expression variability is dominated by partitioning noise (*β* = 0.2 and *ε* = 1 in (20)). (**B**) Comparison of the steady-state protein concentration noise 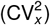 from single-cell (green circles) and population perspectives (brown squares) calculated from simulations of the agent-based model. The solid line represents the analytically-predicted noise level (20). (**C**) Sample concentration trajectories for the high intrinsic noise scenario (*β* = 5 and *ε* = 1 in (20)). Other parameters are taken as 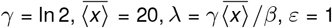, 5000 simulated cells were studied).

- One where the partitioning noise dominates (*β* ≫ *ε*), in which case 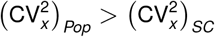 (Figure 4A).
- The other where intrinsic noise dominates (*β* ≫ *ε*), in which case 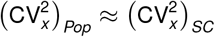 (Figure 4C).

Figure 4B quantifies these differences with increasing intrinsic noise component, i.e., increasing *β*. The key message of this figure is that when intrinsic noise dominates, then both perspectives are similar in terms of the concentration statistics. This can be intuitively understood from our earlier analytical results where both perspectives yield similar protein concentration pdfs (Figure 2B) in the case of perfect partitioning (*ε* = 0). In contrast, the gap between 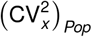 and 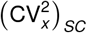 increases as partitioning noise begins to dominate.

## Discussion

In this manuscript, we have investigated stochastic concentration fluctuations in an individual cell over time (single-cell perspective) and across all descendant cells at a fixed time point (population perspective). A key assumption is that the protein of interest is long-lived; thus, its decay is dominated by dilution from cellular growth. We identified two scenarios where the concentration statistics are different between single-cell and population perspectives:

- Expression-growth coupling, where a cell’s proliferation rate depends on the concentration of a specific protein.
- Random partitioning of molecules between daughters during cell division.

Consistent with previous observations [52], our analytical results corroborated with the simulation of agent-based models find an underestimation of noise in the single-cell perspective (Figures 2C & 3C). We discuss these scenarios in more detail below.

Reduction in cellular growth rate with increasing protein concentration was captured phenomenologically through expression (2). This defines feedback in gene expression in which bursts in protein synthesis capture the intrinsic noise in gene expression. The continuous protein dilution between consecutive bursts is defined by the differential equation (3). Our main contribution is to derive exact analytical formulas for steady-state concentration pdfs in both single-cell and population perspectives, as given by (5) and (10), respectively. Before discussing these results, we comment on the no-feedback case. In the absence of any concentration-dilution coupling, both perspectives yield the same gamma-distributed concentration. This result can be generalized to consider transcription feedbacks that are common regulatory features in gene expression [100, 101]. Our analysis in Appendix S9 shows that a constant dilution rate with an arbitrary concentration-dependent burst frequency yields identical concentration distributions in both perspectives.

The presence of dilution-based positive feedback shifts the concentration distribution in the population perspective to lower protein levels (Figure 2B) due to the increased proliferation of low-expressing cells. This explains the lower mean levels and the higher noise and skewness in cell populations compared to the single-cell perspective (Figure 2C). A key qualitative difference is seen in the parameter space where the steady-state concentration distributions exist (Figure 2A). In the single-cell perspective this existence region is defined by the net synthesis rate (*λβ*) being lower than the maximum rate of concentration decrease *γ/k* in (3) that is reached when the protein level is high. The existence region is expanded in the population perspective as cells with higher concentrations proliferate slower, and hence do not contribute significantly to the population. While we have kept the modeling framework simple to obtain analytical insights, models can be refined in the future to consider the scaling of expression rates with the dilution rate [102], explicitly accounting for cell size and cell-cycle effects [103–105], incorporating promoter transcriptional states and mRNA dynamics in more complex gene expression models [106].

In a previous contribution, we investigated dilution-based negative feedback [97], where increasing concentration increases the cellular growth (dilution) rate. This would be the case for many cellular growth factors, where lower concentrations result in lower proliferative capacities [107]. As expected, the results here are opposite to those seen in Figure 2, with the distribution now shifting to a higher concentration in the cell population relative to the single-cell distribution [97]. *In summary, if the expression of a specific gene determines cellular proliferative capacity at an individual-cell level, then the statistical fluctuations in its gene product levels can be qualitatively and quantitatively different between population and single-cell perspectives*.

The discrepancy between single-cell and population perspectives also arises when considering another source of intrinsic noise, the random segregation of molecules between daughters. In the single-cell framework, these random segregation events appear as *jumps* or noise-injections in the concentration at division times (Figure 3B). The statistical properties of these jumps are defined in (16) where the degree of partitioning noise can be tuned through the variable *ε*. We derived the steady-state noise in concentration (20) for an arbitrarily distributed cell-cycle duration and find it to be insensitive to fluctuations in cell-cycle times. In contrast, the corresponding noise within a cell population increases monotonically with increasing randomness in cell-cycle times (Figure 3C). One way to explain this effect is that the randomness in cell cycle times manifests itself as a fluctuation in colony size [108], and larger colonies exhibit higher intracellular concentration fluctuations resulting from the accumulation of noise-injecting division events. Thus, while both perspectives predict similar noise levels for a fixed cell cycle duration, the noise gap increases with 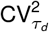, and the noise at the population level is approximately twice the noise in single cells when the cell cycle timing is distributed exponentially (Figure 3C).

A key limitation of our modeling approach is that cell-cycle duration is considered to be *timer*, that is, each duration is independently and identically distributed. It is well known that, if cells grow exponentially in cell size along the cell cycle, then such timer-based models are not able to provide cell size homeostasis, that is, the variance in cell size grows unboundedly over time [109, 110]. We address this limitation by modifying the agent-based model to explicitly consider the size dynamics of individual cells and implemented size control according to *adder* – the size added from cell birth to division is not correlated with the new-born size [111–116]. Statistics computed from simulating these cell-size homeostatic models are presented in Appendix S8 and recapitulate the qualitative finding of Figure 3C: Protein noise in a cell population is more sensitive to fluctuations in added size compared to the single-cell perspective. Interestingly, these simulations show that the single-cell concentration noise that is invariant to 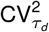 in the timer model (Figure 3C), increases slightly with 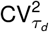 in the adder model (Figure S3). The agent-based models used in this study have been uploaded to *zenodo* for the research community and can be modified to include other types of size control mechanisms and more complex biochemical processes of gene expression [98].

In summary, we have determined differences in studying gene expression in isolated cells versus expanding cell lineages. Although we specifically considered stable proteins, these results can be adapted to other types of biomolecules, such as mRNAs & metabolites, and extended to study intracellular differences in chromosome abundance, plasmids and organelles [117–119]. The focus on intrinsic noise mechanisms (bursting and partitioning noise) can also be generalized to explore extrinsic noise through parametric fluctuations in gene product synthesis/decay rates [120–123]. For example, recent work has reported random fluctuations in translation rates in *Schizosaccharomyces pombe* that dissipate quickly within a cell cycle [124]. Finally, it will be interesting to test our predictions with single-cell expression data using lineage tracking via cellular barcodes, where the extent of cell proliferation can be directly linked to gene expression patterns and their corresponding statistical fluctuations [62].

## Acknowledgments

P.B. and I.Z. have been supported by the Slovak Research and Development Agency under contract No. APVV-18-0308 and the VEGA grant 1/0755/22. A.S. acknowledges support from NIH-NIGMS through grant R35GM148351.

## Author Contribution

I.Z. and P.B. developed the theory for the model of feedback in protein dilution. Z.Z., Z.V. and A.S. proposed the theory for the protein partitioning model. Z.Z. and C.N. developed the stochastic simulations. Z.Z, I.Z. and C.N. designed the plots. I.Z., Z.Z, C.N and A.S. wrote the article. A.S. and P.B. directed the research.

## Appendices

### S1 Appendix. Feedback in dilution. SC model: Chapman–Kolmogorov equation

In the single-cell model we study the dynamics of protein concentration *x* (*t*), which follows the following rules. Protein is produced instantly in portions (bursts) of random size. These bursts arrive at a rate rate *λ* and with a size drawn from an exponential distribution with mean *β*. The protein studied is considered to be relatively stable with a null degradation rate. Then, between the bursts, the protein dilutes exponentially with constant rate *γ*, which corresponds to the cell growth rate. Biologically, this means that the cell cycle duration is distributed following the exponential distribution with mean 1*/γ*. In this appendix, we provide a solution approach for the protein pdf in the single cell model with the feedback in the protein dilution.

The time evolution of the probability density *p*_*SC*_ (*x, t*) is described by the Chapman-Kolmogorov equation:

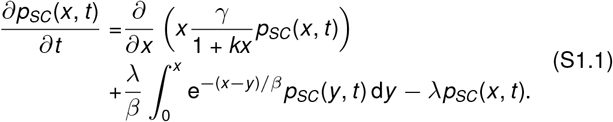

The first term of (S1.1) captures the deterministic drift of probability due to dilution. The last two terms are related to stochastic dynamics. The integral represents the bursts that end at a concentration of *x* and the negative is related to the jumps (burst) when the concentration abandons the state *x*.

To find the stationary distribution, we write it as a probability conservation equation. It is done by collecting the last two terms of (S1.1) as per Leibniz integral rule:

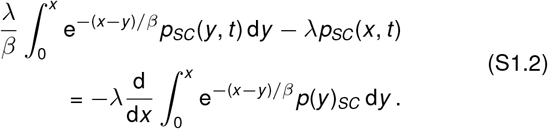

Then, we set *∂p*_*SC*_/*∂t* = 0 in the expression (S1.1), and integrate the result. It yields an integral equation:

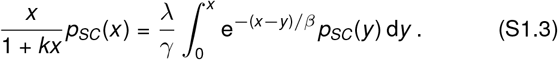

The method to obtain the solution of (S1.3) is based on the Laplace transform. We rearrange that equation in the following way:

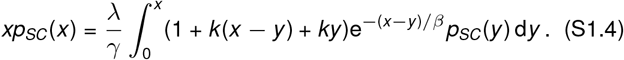

It allows us to represent the right-hand side of equation as the sum of three distinct convolutions:

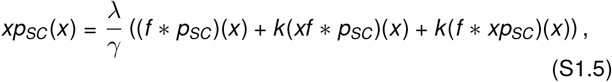

where by *f* we denote the exponential function, that is, *f* (*x*) = e^*−x /β*^. By the asterisk in (S1.5), we denote the convolution of two functions 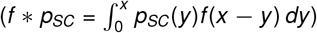, which helps us to solve the problem in the space of Laplace transforms.

As a quick introduction, we define the image *P*(*s*) = ℒ{*p*_*SC*_} (*s*), a function with argument *s*, as the Laplace transform of the function *p*_*SC*_ (*x*) following the relationship:

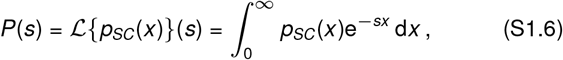

with properties:

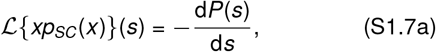

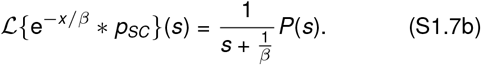

Applying the Laplace transform to (S1.5), it becomes a separable differential equation:

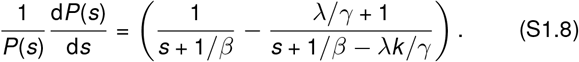

The general solution of (S1.8) is given by:

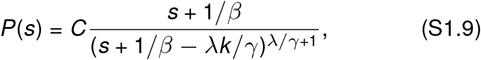

where *C* is an arbitrary constant. Note that the right-hand side is a power function of the Laplace variable *s*, which is shifted by value *η* = 1*/β* −*λk /γ*. In order to return to the original function *p*_*SC*_ (*x*), we use the following relation [125]:

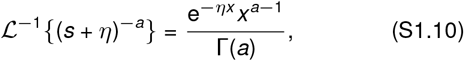

where ℒ^*−*1^ is the inverse Laplace transform to (S1.6). After applying it to (S1.9), we obtain:

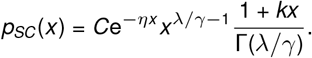

We set *C* so that 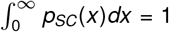 satisfies the normalization condition for the probability density function. This results in the stationary protein distribution:

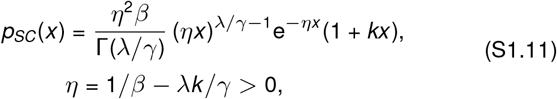

The obtained density (S1.11) is a mixture distribution, thus it can be represented as a weighted sum of two distinct gamma distributions:

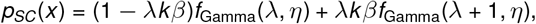

where by *f*_Gamma_ we denoted pdf of Gamma distribution. Note that this distribution is unimodal for any permissible values of parameters *λ, β, γ*, and *k*. This distribution has its peak at zero if *λ/γ <* (4*k β*−1)*/*4(*k β*)^2^ when *k β >* 1*/*4, and if *λ/γ <* 1 when *k β <* 1*/*4; otherwise, the peak is non-zero.

By definition, the *n*-th raw moment is given by the integral 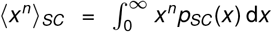; we then use (S1.11) to obtain a closed expression for the *n*-th raw moment:

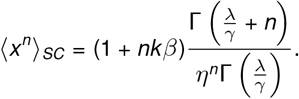

In the absence of the dilution regulation as *k* = 0, *p*_*SC*_ (*x*) becomes the probability density function of the unregulated gene expression *p*(*x*) [67]:

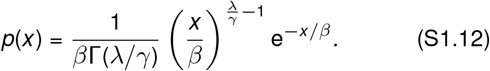

### S2 Appendix. Feedback in dilution. Population model: Population balance equation

In this appendix, we study the feedback in the protein dilution, which is now implemented in the population model and affects not only the protein level but also the proliferation rate. Here, *µ* – the population growth rate – cannot be determined immediately due to the presence of feedback, which affects the cell cycle time and thus the population growth rate. An implicit assumption of the model is that cell volume is strictly increasing at rate *γ/*(1 + *kx*). Consequently, division events are governed by a nonhomogeneous Poisson process with rate *γ/*(1 + *kx*). An appropriate *µ* is required to keep the average cell volume constant, in order to prevent its infinite expansion or diminution to zero.

The expected population density *h*(*x, t*) (number of cells with concentration *x* at time *t*) satisfies the population balance equation:

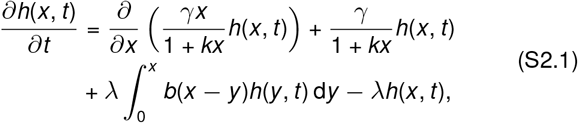

where *b*(*x*) is pdf of the exponential distribution of the burst size.

We start by collapsing the last two terms according to the Leibniz integral rule as per (S1.2). Subsequently, we use the Fourier method, that is, we assume that the population density function can be represented as a separable function:

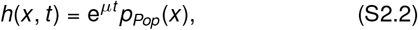

where the principal eigenvalue *µ* gives the population growth rate and the principal eigenvector *p*_*Pop*_(*x*) gives the protein distribution. Then (S2.1) becomes

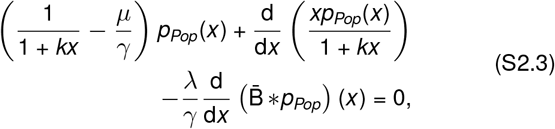

where 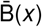 is the ccdf corresponding to *b*(*x*), that is, 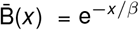.

We define an auxiliary function *q*(*x*) = *p*_*Pop*_(*x*)*/*(1 + *kx*), which we substitute into the equation above; thus we apply the Laplace transform explained in Appendix S1, which gives us the relationship:

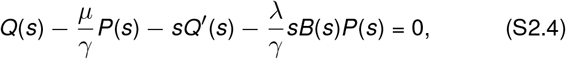

where *P*(*s*), *Q*(*s*), and *B*(*s*) are the Laplace images of functions *p*_*Pop*_(*x*), *q*(*x*), and 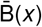 respectively, defined as per (S1.6). Applying the Laplace transform directly to the function *q*(*x*), one can obtain:

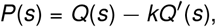

which is used to transform (S2.4) into an ODE for *Q*(*s*):

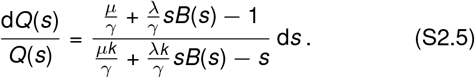

Despite the separable form of this equation, additional complexity is brought about by the generalization of the burst size distribution. However, since 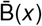 corresponds to the exponential distribution, its Laplace image *B*(*s*) is known:

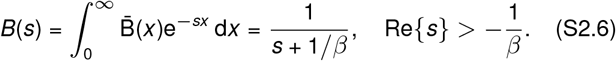

We substitute (S2.6) into (S2.5) and obtain:

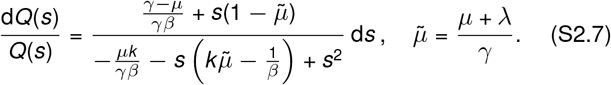

The right-hand side of (S2.7) can be simplified by partial fraction decomposition. The quadratic in the denominator has two roots, which are real and given by

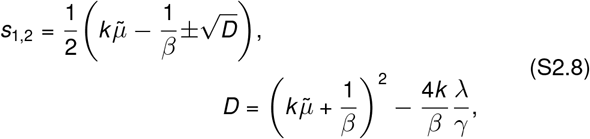

where *s*_1_ *>* 0 and *s*_2_ *<* 0 is for any positive values of *λ, β, γ*, and *k*. The partial fraction decomposition of (S2.7) leads to:

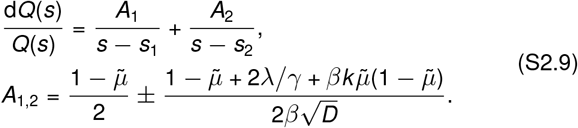

The solution of (S2.7) is

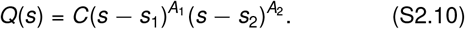

The Laplace transform (S2.10) must be analytic in the complex half-plane Re(*s*) *>* 0, implying that *A*_1_ ∈ {0, 1, 2, …} In particular, the principal eigenvalue is obtained for *A*_1_ = 0; it implies

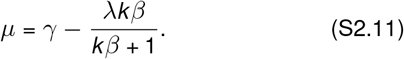

Substituting (S2.11) into (S2.8) and (S2.9), we obtain strictly negative values; we introduce additional variables for their opposite values:

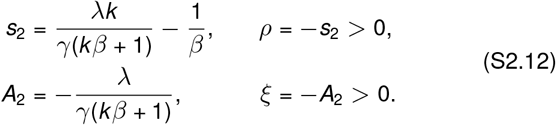

Inserting *s* = 0 into (S2.4) and using the normalization condition *P*(0) = 1 yield *Q*(0) = *µ*, which is used to find the value of *C* in (S2.10). Finally, applying the inverse Laplace transform to (S2.10) and returning to the initial function *p*_*Pop*_(*x*), we obtain:

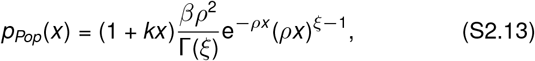

where the constants *ρ* and *ξ* are defined in (S2.12).

**Figure S1:**
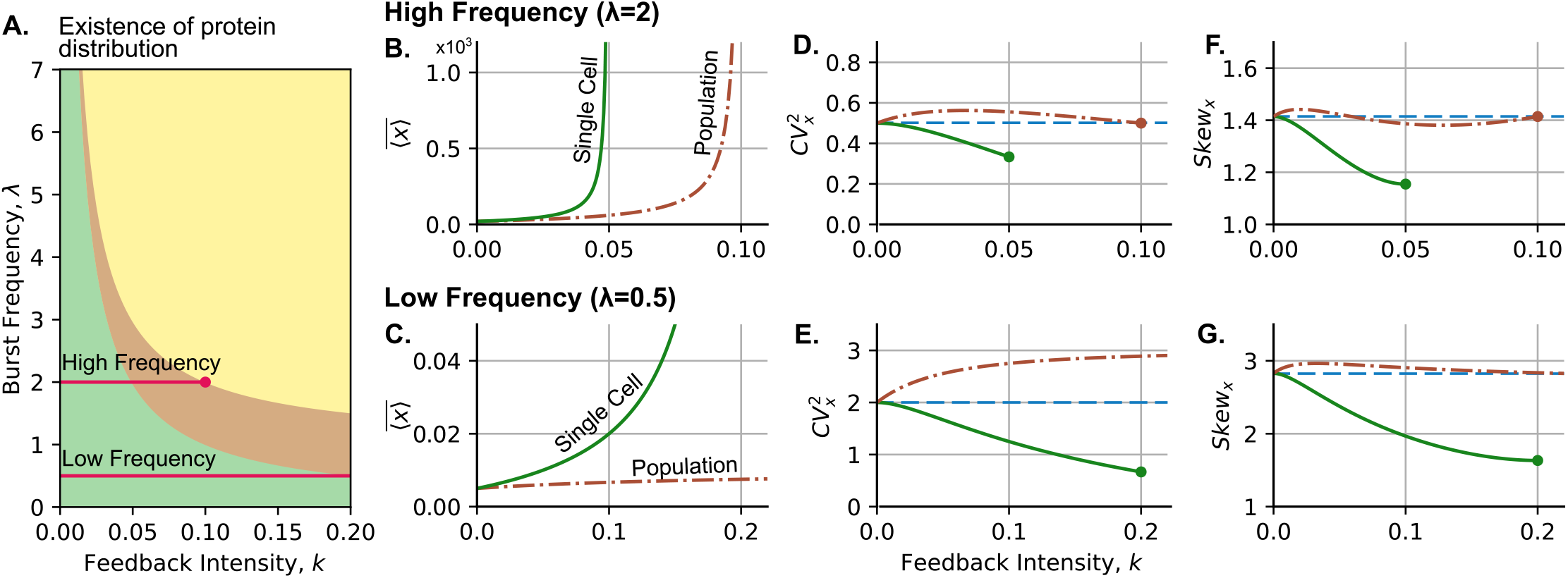
Dilution-based feedbacks result in noiser and more right-skewed concentrations across cell populations as compared to the single-cell perspective. (**A**) Phase diagram of distribution existence similar to Figure 2A. Red line represents increase of *k* keeping *λ* = 2 and the low frequency *λ* = 0.5. Mean protein concentration: (**B**) for high frequency and (**C**) for low frequency. Protein noise: (**D**) for high frequency and (**E**) for low frequency. Protein Asymmetry: (**F**) for high frequency and (**G**) for low frequency. On (**B**)–(**G**) are shown statistics of single cell (green solid line) and population (brown dash-dotted line) compared to the unregulated case (blue horizontal dashed lines). Parameters: *β* = 10, *γ* = 1.

As well as (S1.11), stationary protein distribution *p*_*Pop*_(*x*) is a mixed distribution, i.e.,

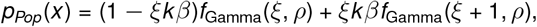

which is also unimodal for any set of parameters *λ, β, γ*, and *k* satisfying *ρ >* 0.

Note that without regulation, as *k* = 0, the pdf of the protein concentration in the population model is identical to the one in the single cell (S1.12). Finally, we obtain moment expressions for the *n*-th raw moment:

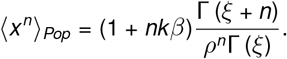

### S3 Appendix. Moments analysis of the effects of the feedback strength keeping the burst frequency constant

In the main text, we focus on asymptotic behavior in the case when the burst frequency is adjusted so that the mean concentration remains fixed (7). In this appendix, we study protein noise and skewness as functions of feedback strength *k*, with parameters *λ/γ* and *β*.

In the single cell, the protein distribution *p*_*SC*_ (*x*) (5) has the existence condition *η >* 0, from which it follows that *k* must be within the interval 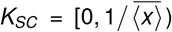, with 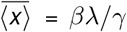 for a given set of parameters (*λ, β, γ*). It is clear that as *k* approaches 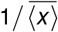, in the expression (6a), the mean value 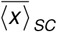 diverges due to a singularity in the denominator (Fig. S1B, green line). To explore the behavior of the noise level (6b) and skewness (6c) on *K*_*SC*_, we use the first derivative test. For the noise level, we find that 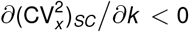 meaning that 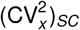 is decreasing function of *k* within *K*_*SC*_ (Fig. S1D, green line). We also use this approach for the skewness; its derivative is given by:

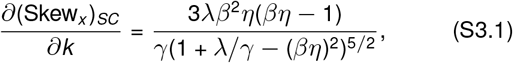

where the quadratic function in denominator is always positive (it is concave and its two zeroes are not in *K*_*SC*_), then *∂*(Skew_*x*_)_*SC*_ */ ∂k <* 0 for any *k* ∈ *K*_*SC*_. We find that both 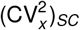 and (Skew_*x*_)_*SC*_ are monotonically decreasing functions with local maxima and minima being left and right endpoints of the interval *K*_*SC*_, respectively. In particular:

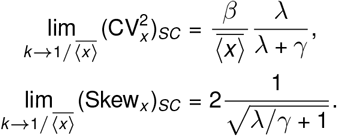

In conclusion, for a given production flow *λβ*, the feedback of any strength reduces protein noise at the single cell level (green lines in Figs. S1D–S1E) and makes the distribution less skewed (green lines in Figs. S1F–S1G) compared to the unregulated expression (blue horizontal lines in corresponding figures).

We follow a similar approach for the statistics of from the population perspective, where the permissible interval of *k* is 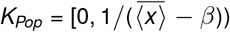, for *λ > γ* (the high frequency limit), and *K*_*Pop*_ = [0, ∞), for *λ < γ* (the low frequency limit). The behavior of mean, noise level, and skewness depends on the ratio *λ/γ*. In a low-frequency mode (*λ < γ*), we obtain:

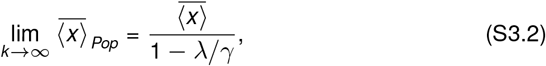

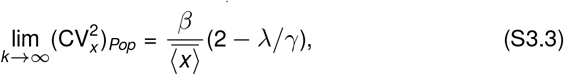

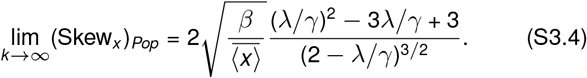

Then the protein distribution in the population with the low transcriptional frequency (*λ < γ*) has higher, but always bounded statistics compared to unregulated case. This behaviour is shown in the second row of Fig. S1.

In high frequency mode (*λ > γ*) as *k* reaches the right endpoint of *K*_*Pop*_, the mean value 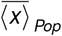 diverges and the statistics become identical to the unregulated case:

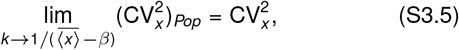

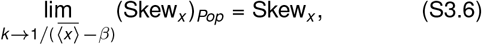

which is shown in the first row of Fig. S1.

The first derivative of the squared coefficient of variation,

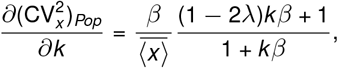

has a single root at 1*/*(2*λ*−1)*β*, indicating that over the interval 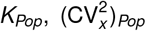 is monotonically increasing if 2*λ < γ*; otherwise, it is concave (Figs. S1D–S1E). The first derivative test for skewness involves analysis of a cubic equation, which was done numerically. We conclude that the low frequency leads to (Skew_*x*_)_*Pop*_ *>* Skew_*x*_ on the whole interval *K*_*Pop*_, maximum is reached within *K*_*SC*_ (Fig. S1G). The high frequency leads to minor fluctuations of (Skew_*x*_)_*Pop*_ around Skew_*x*_ with single intersection within *K*_*Pop*_ (Fig. S1F).

Overall, we use the statistics of the unregulated case as a critical point for comparison of both perspectives. We conclude that for given production rate *λβ* and admissible values of *k >* 0 protein distribution in the population is always noisier and more right-skewed compared to the single-cell one.

### S4 Appendix. Statistical properties of protein partitioning

In this appendix, we derive the statistical properties of the jumps that the protein concentration performs during partitioning.

#### S4.1 Statistics of protein level during partitioning

First, consider a scenario in which just before cell division, the protein concentration within the cell is indicated as 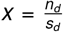, where *n*_*d*_ represents the number of protein molecules and *s*_*d*_ is the cell size. As the cell divides, its size is halved, and each daughter cell inherits a size of 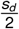. The total number of molecules, *n*_*d*_, is assumed to segregate with a probability of 0.5 for each daughter. Consequently, the number of molecules in each daughter cell follows a binomial distribution with parameters *n*_*d*_ and 0.5.

Given these premises, the mean and variance of the protein count in each daughter cell, denoted as *n*_+_, have the moments of the binomial distribution:

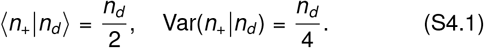

This allows us to find the mean and variance of the protein concentration in a daughter cell:

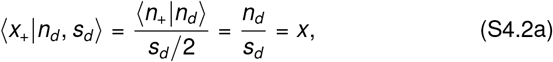

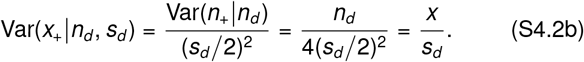

In the general case, protein partitioning may not be perfectly binomial and have complex mechanisms. To generalize expression (S4.2b), let us define the statistics for the protein level after partitioning:

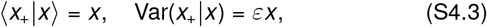

where *ε >* 0 is a constant that reflects these effects of size scaling and complex partitioning. For the case of binomial partitioning, *ε* = 1*/s*_*d*_.

#### S4.2 Simulation of protein partitioning

During partitioning, protein levels change following the statistics shown in (S4.3) with the additional constraint that 0 *< x*_+_ *<* 2*x*. In our simulations, we model the protein partitioning, proposing that the protein level after division follows:

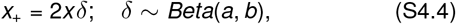

which means that we multiply the protein before division *x* by two times a random variable *δ* which is beta-distributed with shape parameters *a* and *b*. To obtain the desired stochastic properties of *x*_+_ as defined in (S4.3), we have to satisfy ⟨*δ*⟩ = 1*/*2 and Var(*δ*) = *εx*. Using these constrains, we obtain the values for the shape parameters of the beta distribution (S4.4):

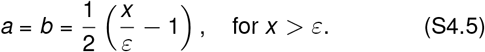

Due to the properties of the beta distribution, the limit *x* = *ε* means that *δ* takes the values of 0 and 1, each with equal probability. This scenario represents the highest possible variance for any distribution confined to the interval (0,1). For a biological interpretation, if *ε* satisfies *ε* = 1*/s*_*d*_, then *x* ≤ *ε* means that the progenitor cell either does not contain any protein molecules (leaving nothing to segregate) or contains exactly one. Hence, during cell division, this single molecule is inherited randomly by one of the descendants, leaving the other cell without any molecules. In our simulations, when *x* ≤*ε*, one descendant cell receives a protein concentration of 2*x*, while its counterpart receives none.

### S5 Appendix. Derivation of the protein steady-state mean and noise: single cell perspective

In this appendix we present our theoretical framework for calculating the statistical moments of protein concentration, taking into account the role of molecular partitioning in gene expression. For this purpose, we use the framework of time-triggered stochastic hybrid systems (TTSHS) [126–128], which integrates the continuous dynamics of protein dilution with two families of resets: (i) cell division and (ii) protein synthesis in bursts.

To model cell division we leverage the theory of renewal processes that is a generalization of the classical Poisson process. Here the time between two successive events is an independent and identically distributed random variable following an arbitrary distribution 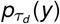, i.e., the probability

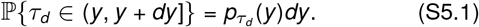

To model division events we introduce a timer *τ* that is set to (*τ* = 0) when a cell is born, and increases linearly with time along the cell cycle

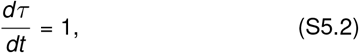

together with decay in protein concentration

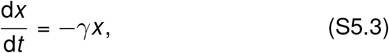

as a per fixed dilution rate *γ*. Cell division events occur probabilistically with propensity (or hazard rate)

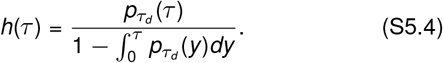

More specifically, given the state of the timer *τ*, the probability of a division event occurring in the next infinitesimal time interval (*t, t* + *dt*] is *h*(*τ*)*dt*. Whenever this division event occurs, the timer and concentration are reset as

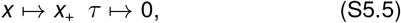

with *x*_+_ representing the protein concentration in one of the randomly-chosen daughters in the single-cell perspective. The statistical properties of *x*_+_ were described in Section S4.1. The case of Poisson process is recovered if 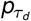 is precisely an exponential distribution with mean ⟨*τ*_*d*_⟩, then as per (S5.4) the cell division propensity *h*(*τ*) = 1*/* ⟨*τ*_*d*_⟩ would be a constant. Note that in this model formulation *τ* is a stochastic process and the steady-state expected value of any generalised hazard-rate satisfies:

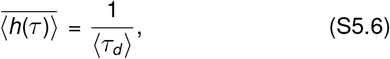

as shown in [127]. Having explained the timing of cell division, we next describe protein synthesis events occurring in stochastic bursts. Synthesis events occur as per a constant propensity *λ*, and each event increases the protein concentration as per the reset

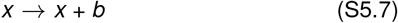

where *b* an independent and identically distributed random variable following an exponential distribution with mean 1*/β*. This model of protein synthesis can be further generalized where both the time interval between two successive bursts events and the burst size *b* follow arbitrary distributions [77]. In summary, the deterministic dynamics (S5.2)-(S5.3) together with cell division and protein bursting events with corresponding resets (S5.5) and (S5.7), respectively, define a time-triggered stochastic hybrid systems (TTSHS) which is itself a special class of PDMPs.

We refer the reader to [126, 127] for details on deriving time evolution of the statistical moments of TTSHS state space. Using the statistical properties of *x*_+_ in (S4.3), the time evolution of the first- and second-order moments of the protein concentration follow the system of differential equations

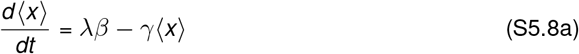

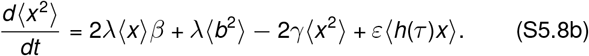

Using the fact that at steady-state (see Theorem 1 in [127])

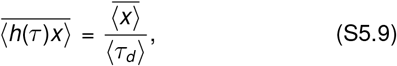

and ⟨*b*^2^⟩ = 2*β*^2^, *γ*⟨*τ*_*d*_ ⟩ = ln 2, solving (S5.8) at steady state yields

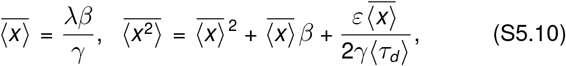

which results in the following expression for concentration noise

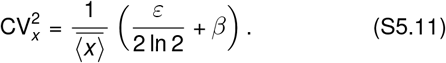

### S6 Appendix. Partitioning noise with deterministic cell cycle duration

In this appendix, we solve the moments of protein concentration from the single-cell and population perspectives when the cell cycle has a deterministic duration. As a final result, we show that the protein concentration noise is the same in both perspectives.

**Figure S2:**
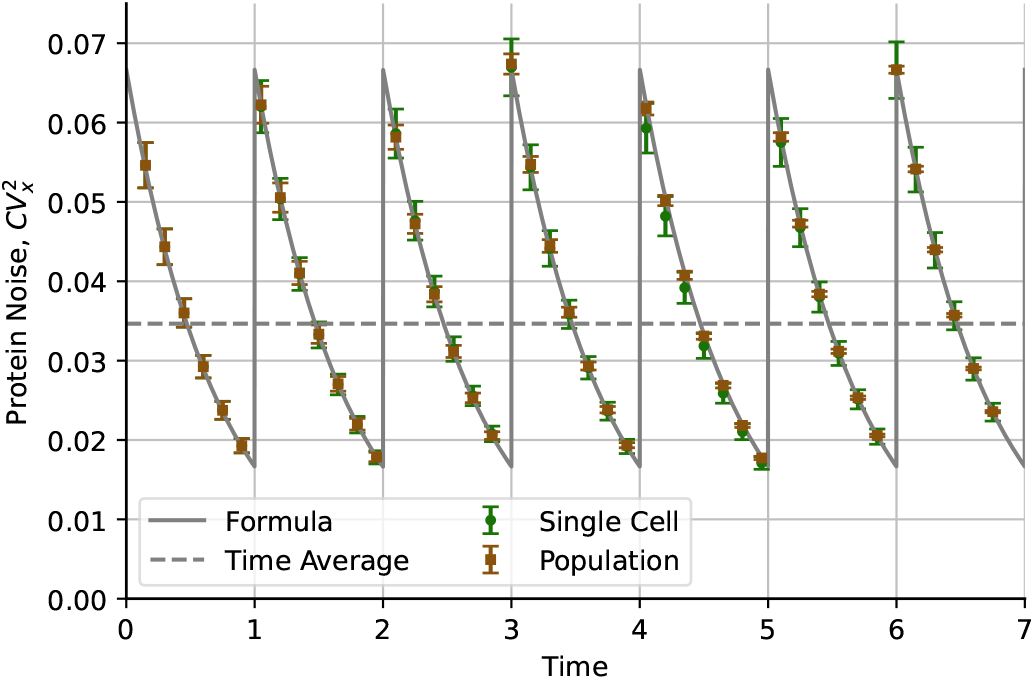
Protein concentration noise in single-cell and population perspectives follows the same dynamics for fixed cellcycle duration. The line represents the protein noise calculated using (S6.11). The scatter-plot shows the results of simulations (green circles: single-cell, brown squares: population). Parameters: *γ* = ln 2, *λβ* = 20*γ, ε* = 1 *τ*_*d*_ = 1, 5000 simulation replicas.

#### S6.1 Noise at single cell level

First, we solve the moment dynamics for the single-cell perspective. We neglect protein bursting and therefore assume that protein concentration evolves deterministically as per (15). In this way, protein concentration trajectories consist of continuous dynamics interrupted by random-size jumps that represent partitioning during division. These divisions occur with a period of deterministic duration *τ*_*d*_ and therefore we can solve the ODE for protein concentration (15) applying periodic boundary conditions. This means that the system has the same properties after each multiple of *τ*_*d*_.

We define the timer *τ* to track the time since the previous division following 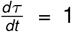. During the cell cycle *τ* ∈ [0, *τ*_*d*_), the protein concentration evolves as per (15) which has the solution:

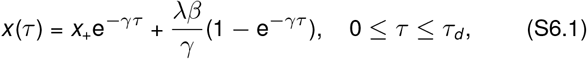

where we have used the property (S5.5) that, during division, protein levels jump from *x* (*τ*_*d*_) to the random variable *x*_+_ with unknown moments ⟨*x*_+_⟩ and 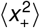. Using the periodic boundary conditions, and considering that during division the mean concentration does not change, we have:

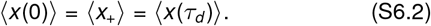

Taking the expected value of (S6.1) and using the boundary conditions (S6.2), we solve for ⟨*x*_+_⟩:

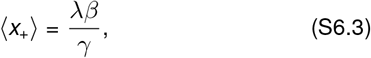

which also solves the mean concentration as function of *τ* :

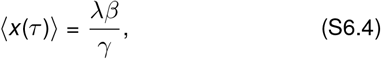

which is consistent with the fact that the division does not change the mean concentration level. To obtain the variance of *x* over time, we find the second moment ⟨*x*^2^(*τ*)⟩. Since the protein trajectory for given *x*_+_ is deterministic, the square of the protein level follows:

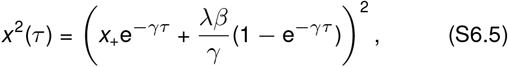

which has the expected value given *τ* :

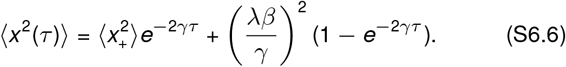

Next step is to apply the boundary conditions. During cell division, this is, when time is a multiple of *τ*_*d*_, protein level jumps from *x* (*τ*_*d*_) to *x*_+_ and the variance increases as:

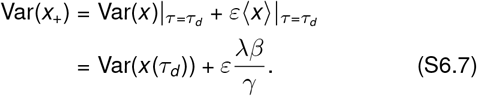

Notice that Var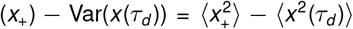 because the averages are identical 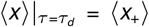. Therefore, we conclude that:

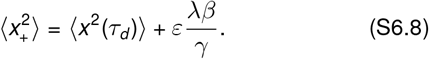

It is possible to replace ⟨*x*^2^(*τ*_*d*_)⟩ from equation (S6.6) into (S6.8) and solve for 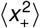 to obtain:

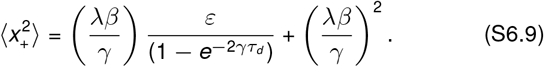

Substitution of this second moment of *x*_+_ into (S6.6) yields:

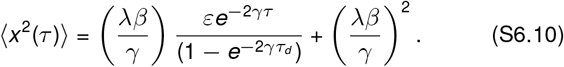

Using also ⟨*x* (*τ*) ⟩, it is possible to obtain the protein noise throughout the cell cycle:

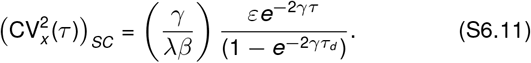

Figure S2 presents the trajectory of 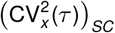 using (S6.11); the comparison with simulations shows a relatively good match.

#### S6.2 Noise at population level

In this subsection, we demonstrate that the noise at the population level coincides with the noise at the single-cell perspective. From the population perspective, after division at *τ* = 0^+^, we continue to follow both daughter cells. While one daughter cell inherits the protein level *x*_+_, the other inherits the level 2 ⟨*x* (*τ*_*d*_)⟩ − *x*_+_. This is because the mean concentration of both cells just after the division the same before the division, that is ⟨*x* (*τ*_*d*_)⟩. After a timer *τ* ∈ [0, *τ*_*d*_), the protein concentration in the first daughter is presented in (S6.1). The second daughter follows the dynamics:

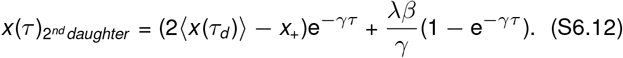

Using the property ⟨*x*_+_⟩ = ⟨ *x* (*τ*_*d*_) ⟩, we obtain that the mean concentration for both daughters at equilibrium are identical:

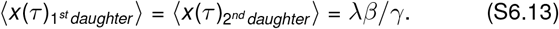

To obtain the dynamics of the second-order moment for the second daughter, we take the square of the equation (S6.12) and take the average:

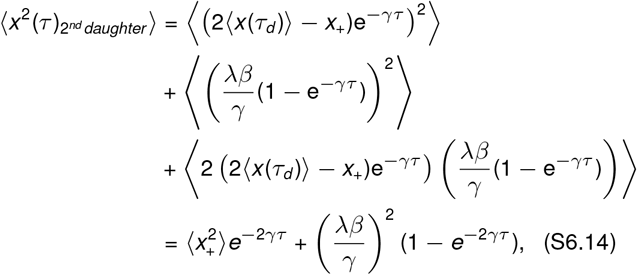

where we have used 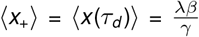. Notice that equation (S6.14) is the same as (S6.6) which has solution (S6.10):

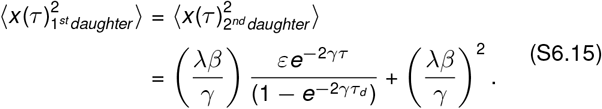

The first and second order moments of both daughters are identical, so that the 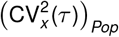 during the cell cycle *τ* ∈ [0, *τ*_*d*_) can be obtained as follows:

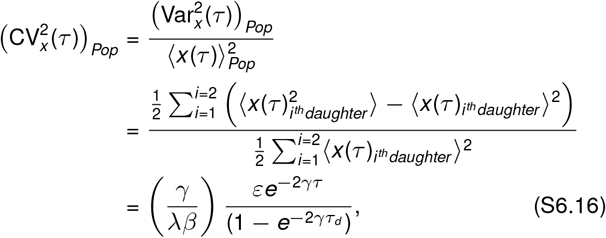

which is the same as the single cell noise given by (S6.11). Consequently, it can be shown that after the *n*^*th*^ division, the noise level within the population is equal to that of single cell, i.e.,

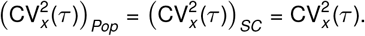

This result is verified using the simulations, which is shown in Figure S2. It can be shown that with the parameters *γ* = ln 2*/τ*_*d*_ and *τ*_*d*_ = 1, the time-averaged 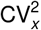 is given by,

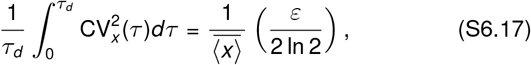

which corresponds to the dashed horizontal line in Figure S2.

### S7 Appendix. Simulation Algorithm for estimating the protein statistics in SC and population model

In this appendix, we present the basis of the simulation algorithms used in the main article. The main idea of Algorithm 1 is to produce two time intervals for the next reactions and select the minimum. The reaction related to that minimum time will be chosen to happen. The first reaction corresponds to the protein burst, and the second is the division event. The main difficulty in the simulation relative to other standard methods such as the classical Gillespie’s algorithm [129] is that the division time is not exponentially distributed, and therefore division is not a memoryless process. This means that each cell has to track the time left for division. In addition, when the population perspective is considered, the number of cells in the population increases during each division. To address this, we designed an agent-based algorithm that models protein levels within each cell over time.

The algorithm has the goal of estimating the protein level *x*^*i*^ of cells in the population. Here, the superscript *i* represents the *i* -th cell of the population. When the **single-cell perspective** is considered, only one descendant is randomly chosen to inherit a beta-distributed protein level during each division, so the population corresponds to one cell. Otherwise, in **population model**, a new cell is added to the population after each division, the number of cells in population grows each division.

The process starts with setting the initial conditions. For each replica, we start the colony progenitor (*i* = 1) with a given protein level *x*^1^. Simultaneously, we set the maximum time to end the simulation *T*. Each cell has two additional variables: the time to the next burst 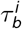 and the time to division 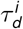. While 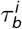 is exponentially distributed with mean 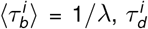 is considered as gamma distributed with shape and scale parameters selected, such as this variable has mean ⟨*τ*_*d*_⟩ and variability 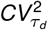. In some cases we consider gene expression as non-bursty, this means that the burst does not occur. This can be done considering 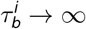 and choosing always the division.

During each iteration, the minimum time *τ*_*min*_ is selected among all 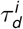 and 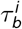, and the respective reaction occurs in the *i* -th cell. Before the reaction, all 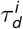 and 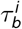 decrease by *τ*_*min*_. Simultaneously, the protein level of all cells evolve following the differential equation:

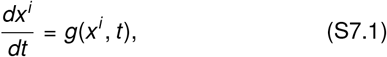

where the function *g*(*x, t*) depends on whether the gene expression is bursty or not. For a bursty protein synthesis the protein dilutes as *g*(*x, t*) = −*γx*. In non-bursty case, protein evolves as *g*(*x, t*) = *λβ* − *γx*, and we reassign 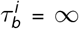 in Algorithm 1, so that the division is the only possible event.

We denote by *n* the cell that is the one with the corresponding minimum time. If 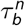 is that minimum, the protein burst is set to happen; otherwise, the division occurs. During the burst, the protein level increases by a jump with size drawn from an exponential distribution with mean *β*. During division, the protein level is reset to a new variable *x*_+_ with mean ⟨*x*_+_⟩ = *x* and variance Var(*x*) = *εx* as explained in Section S4.2.

#### Algorithm 1

Given the initial protein concentration *x*_1_, and the time to complete the simulation *T*, we obtain the end of the simulation the array of protein concentrations at 𝕏 = {*x*^1^, · · ·, *x*^*N*^} for the *N* cells. If the simulation is done for the population perspective, the algorithm saves a new descendant cell every time a cell in the population divides; for the single-cell perspective all descendants are ignored.

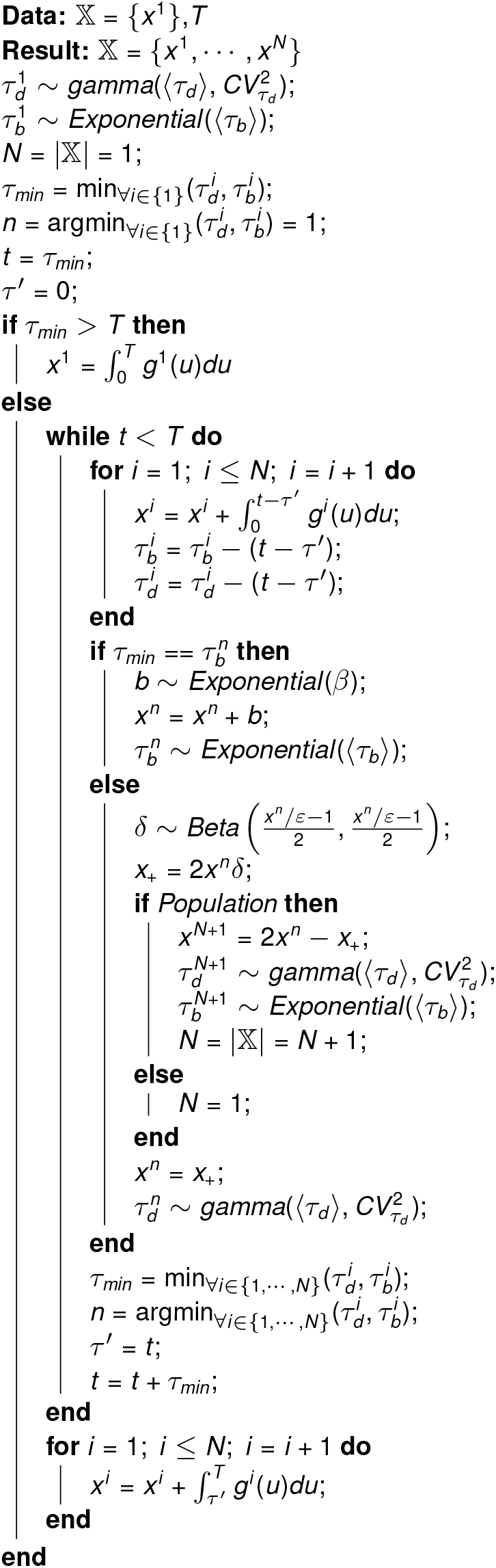

**Figure S3:**
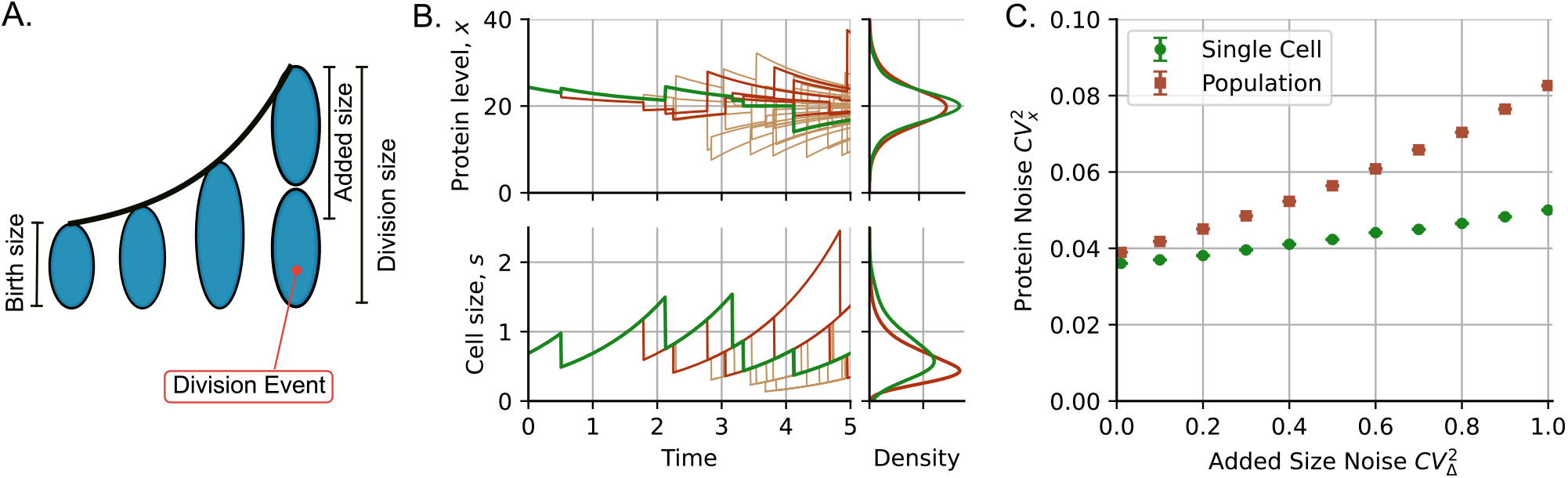
Adder-based cell size homeostasis causes differences in concentration noise levels between the population and single-cell frameworks. (**A**) Cell size dynamics following adder principle. After division, a cell has a given birth size, grows exponentially, and divides once it has expanded by added size. The added size is an independent random variable with a given noise 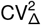. (**B**) Trajectories of protein level (top) and cell size (bottom) in a sample population. Green lines represent the protein levels of a single cell, dark brown are the trajectories of the descendant cells and other trajectories correspond to remaining cells in the colony. (**C**) Comparison of noise in protein levels as a function of noise in added size 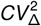. Green scatter plots show single-cell statistics and brown ones are for population. While the noise at the single-cell level changes weakly with the noise in added size, there is a greater change at the population level. Parameters of protein concentration: *λβ* = 20, *γ* = ln(2), *ε* = 1. Parameters of cell size: ⟨*s*_*b*_ ⟩ = 0.5, ⟨*s*_*d*_ ⟩ = 1, Δ is drawn from gamma distribution with ⟨Δ⟩ = 0.5. Single-cell statistics are based on the simulation of 50000 individual cells, population statistics are derived from 1000 colonies after approximately six divisions.

When the population perspective is chosen, during division, a new cell is added to the population. While, randomly, one cell inherits a protein level *x*_+_ with the statistics explained before, the other cells inherit a level 2*x* − *x*_+_. This ensures that the mean protein level across both daughter cells is maintained at *x*. For single cell model, only one of the descendants is randomly chosen to inherit a protein level *x*_+_, maintaining population size constant at one cell.

This simulation is performed across multiple populations, involving 5000 replicates. To estimate the moments of the distribution, for the single cell perspective, we calculate the average across 5000 cells, one from each colony, at the end of the simulation. For the population perspective, statistical estimates are derived from all cells across 5000 colonies, without distinguishing between lineages.

### S8 Appendix. Pop and SC difference for the adder division

We consider that cell size increases exponentially over time with divisions following the *adder* strategy [112] – cells divide after adding, on average, a fixed cell size called *added size*. This added size is presented in Figure S3A as the difference between the cell size at birth and the cell size at division.

During the cell cycle the cell size *s*(*t*) grows exponentially:

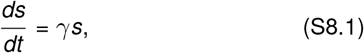

with the same growth rate *γ* as the protein dilution rate. The protein concentration evolves deterministically according to (15) with constant synthesis and dilution rates. During each cell division, the protein partitioning follows the method presented in Appendix S4. Then, according to (S4.3), if the cell has concentration *x* and size *s*_*d*_ at the end of its life cycle, the mean protein concentration in the new cells equals to *x*, with variance *x /s*_*d*_. In Figure S3B, we show the cell protein concentration dynamics (top) and the cell size dynamics (bottom).

During a cell cycle, cells add a size denoted by Δ. For simulations, we assume that Δ is a random independent variable that follows a gamma distribution with mean ⟨Δ⟩ = 0.5 and noise 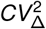. This choice of ⟨Δ⟩ is set so that the mean size at division ⟨*s*_*d*_⟩ = 1, making the results of *adder* comparable to those of *timer* in the main text, following *ε* = 1*/* ⟨*s*_*d*_⟩ = 1.

To obtain the cell cycle duration *τ*_*d*_ for each cell, we register its birth size *s*_*b*_, and then the size at the end of that cycle is *s*_*b*_ + Δ. Given the exponential cell growth, we have 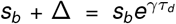, which we use to obtain the cell cycle duration as follows:

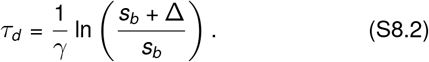

Note that the duration of the cell cycle *τ*_*d*_ is now a variable that depends on *s*_*b*_. This means that, unlike the *timer* approach considered in the text, *τ*_*d*_ is no longer an independent variable.

For simulation, each cell *x*^*i*^ in a colony has parameters: protein level *x*_*i*_, cell size *s*_*i*_, added size Δ^*i*^, time for next burst 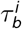 drawn from the exponential distribution, and cell cycle duration 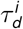 estimated using (S8.2). Then Algorithm 2 shows that based on these parameters each cell colony evolves as follows. The set of all 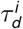 and 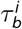 over the cells in the colony defines all possible reaction times. During each iteration, we select the minimum of those times, perform the associated reaction, and allow the system to evolve during that minimum time. The evolution of the colony includes the exponential growth of cells at a rate *γ* with the subsequent dilution of proteins at the same rate. If the selected reaction corresponds to a burst, the protein level of the cell increases by an exponentially distributed quantity with mean *β*. If the reaction is a division event, the protein level is perturbed, as explained in Appendix S4, and, for population level, another cell is added to the colony. The iteration continues until the maximum time *T* is reached.

#### Algorithm 2: Simulation algorithm for cell proliferation following the *adder* division strategy

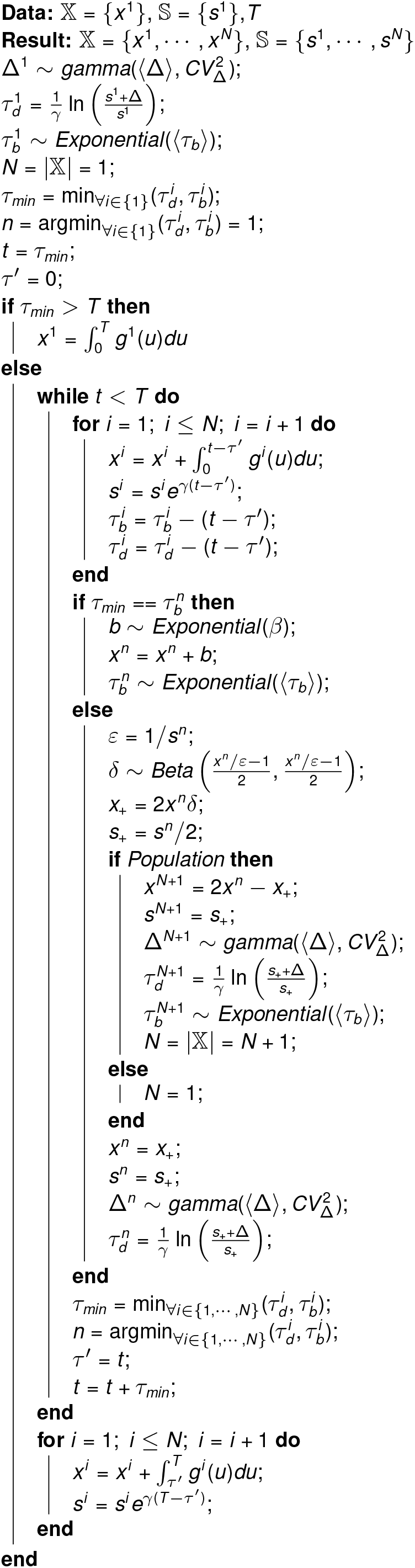

Given that there is no simple way to measure the noise in *τ*_*d*_ because the duration of the cycle is defined by Δ instead, we simulate 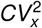 as a function of the noise in the added cell size 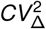. A comparison of the level of noise protein 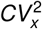 in the population and the single-cell perspective is presented in Figure S3C. We observe that with the *adder* method of cell division, the difference in protein noise level between the population and single cell is similar to the scenario where cell division follows the *timer* strategy (Figure 3C). However, protein noise for a single cell has a slight increase with 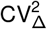. This is because the variance of the protein level after partitioning is a function of *s*_*d*_ and the noise in this variable increases with 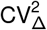.

### S9 Appendix. Feedback in frequency. Identical protein distribution in SC and population model: Proof

When the we generalize the burst frequency to a function *λ*(*x*) of the protein concentration, the time-evolution of the protein concentration pdf *p*_*SC*_ (*x, t*) is governed by the Chapman-Kolmogorov equation:

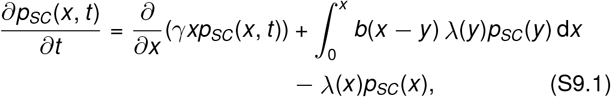

where *b*(*x*) = e^*−x /β*^*/β* is probability density function of the exponential distribution.

To find the stationary distribution, we write it as a probability conservation equation. It is done by gathering the last two terms of (S9.1) as per Leibniz integral rule into the following derivative:

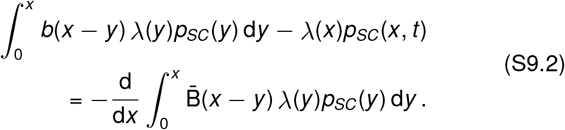

where 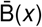 is the complementary cumulative distribution function (ccdf) corresponding to *b*(*x*), i.e., 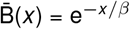.

Finally, the steady state of the system implies that distribution does not change over time, i.e., *∂p*_*SC*_ (*x, t*)*/∂t* = 0. We obtain:

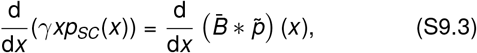

where an auxiliary function 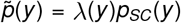 is used for convenient representation of the convolution.

We proceed with the model of the population, where the life-cycle of each cell is identical to one described at the beginning of the chapter. Its composition is given by the function *h*(*x, t*) – number of cells with given concentration *x* at the time *t*. The time evolution of *h*(*x, t*) is described by the population balance equation:

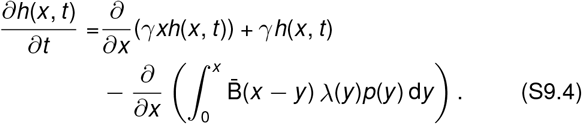

which differs from the Chapman-Kolmogorov equation (S9.1) by the inclusion of the population growth term *γh*(*x, t*).

However, in the absence of the feedback in dilution, we have exponentially distributed lifespans. In this case, the population growth rate depends only on the division frequency, i.e., the eigenvalue *µ* = *γ*. By substitution, we can prove that solutions of PBE satisfy

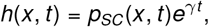

where *p*_*SC*_ (*x, t*) solves the master equation (S9.1). From it follows, in particular, that

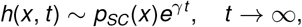

where *p*_*SC*_ (*x*) solves the stationary problem for the single-cell perspective (S9.3). Then the stationary distributions of the protein concentration for single-cell perspective *p*_*SC*_ (*x*) and population perspective *p*_*Pop*_(*x*) are identical.

